# Waiting times in a branching process model of colorectal cancer initiation

**DOI:** 10.1101/2022.06.22.497226

**Authors:** Ruibo Zhang, Obinna A. Ukogu, Ivana Bozic

## Abstract

We study a multi-type branching process model for the development of colorectal cancer from initially healthy tissue. The model incorporates a complex sequence of driver gene alterations, some of which result in immediate growth advantage, while others have initially neutral effects. We derive analytic estimates for the sizes of premalignant subpopulations, and use these results to compute the waiting time distributions of novel driver mutations. The approach presented here is relevant to the study of finite-time properties of multi-type branching process models in which growth rates of consecutive types are non-decreasing.

## 1. Introduction

Following the seminal work by Armitage and Doll in the 1950s [1, 2], and Knudson [3] in the 1970s, cancer initiation has been identified as a multi-step process, whereby cells sequentially accumulate driver mutations required for malignant transformation. These works investigated age-dependent incidence rates for human cancers and argued that a multi-step model best explains the data across different cancers. Since the advent of genome sequencing, the molecular mechanisms of many cancer driver mutations have been described [4–8].

Multi-type branching processes, with types corresponding to genotypes or cell states [9, 10], have emerged as a viable model for studying cancer evolution. Multiple aspects of cancer evolution have been modeled, including initiation [11, 12], progression [10, 13–15], metastasis [16–18], and resistance to therapy [19–23]. Evolutionary dynamics are closely related to the relative fitness advantages conferred by mutations. Durrett and Moseley [10] analyzed a scenario in which the clonal growth rate strictly increases after each mutation, and computed distributions for clonal sizes and waiting times. Nicholson and Antal [23] studied a general framework wherein wild-type individuals have the largest fitness (growth rate), which could be applied to cases involving drug resistance. Random fitness advantages have also been investigated [24, 25].

Colorectal cancer (CRC) is one of the most common cancers in the United States [26]. It has been shown that mutations in just three driver genes, usually inactivation of two tumor suppressor genes and activation of an oncogene, are sufficient to trigger the initiation of CRC [11, 27, 28]. Typically, this involves a total of five genetic alterations, as inactivation of a tumor suppressor gene (TSG) requires inactivation of both alleles, and activation of an oncogene only requires a mutation in one allele of the gene [29]. Recent work [11] studied a five-step branching process model for the initiation of colorectal cancer that involves the three most commonly mutated driver genes in colorectal cancer: tumor suppressors *APC* and *TP53* and oncogene *KRAS.* The study found that, in the majority of cases, the driver mutations accrue in a specific order, with inactivation of *APC* followed by activation of *KRAS* and inactivation of the *TP53* gene.

Following [11], we study the single most likely mutational pathway to colorectal cancer. We model the dynamics using a multi-type branching process that starts from *N* wild-type crypts, small tubular assemblies of cells that line the intestinal epithelium [30, 31]. As the process evolves, individual crypts stochastically obtain driver mutations, with mutation rates determined by the genotype of the crypt and the driver gene in question. We allow growth rates of consecutive subpopulations to be equal, which leads to considerable mathematical difficulty. Furthermore, instead of starting with a growing population initiated by a single cell, we consider the case where the process is initiated from a large population of nondividing wild-type crypts. Using results from the theory of martingales, we compute the waiting time distributions associated with each step of the process. Finally, we compare analytic results for the waiting time distributions to those from exact computer simulations of the process.

The approach presented here can be extended to other multi-type branching process models in which the growth rates of subsequent types are non-decreasing. For colorectal cancer, one can use a similar approach to compute the waiting time distributions for other mutational pathways. The quantitative estimates provided here may also help better understand the lifetime risk of CRC.

## 2. Model

Let *N_i_* (*t*) be the stochastic process that counts the population of type-*i* crypts at time *t*. In a single step, a type-*i* crypt transforms into a type-(*i* + 1) crypt through either mutation or loss of heterozygosity (LOH) at rate *u_i_*. Here we consider the most common mutational pathway observed (see Table 1) over the course of an 80-year human lifespan, i.e. 0 ≤ *t* ≤ 80. We assume that independently from mutations, type-*i* crypts follow a pure birth process with rate λ_*i*_. The growth rate of a crypt is determined by its genotype. For wild-type crypts, divisions are very rare [32] so we set their growth rate to zero. It has been shown that inactivation of *APC* and/or activation of *KRAS* provides a fitness advantage to mutated crypts, leading to clonal expansion [33, 34]. In contrast, under normal conditions *TP53* inactivation alone does not provide a fitness advantage [30]. These findings are reflected in the choice of growth parameters in our model (see Table 2).

**Table 1:** Most common pathway to CRC initiation.

| Step | Rate | Biological process |
| --- | --- | --- |
| $N_0 \rightarrow N_1$ | $u_1$ | *APC* LOH |
| $N_1 \rightarrow N_2$ | $u_2$ | *APC* mutation |
| $N_2 \rightarrow N_3$ | $u_3$ | *KRAS* mutation |
| $N_3 \rightarrow N_4$ | $u_4$ | *TP53* LOH |
| $N_4 \rightarrow N_5$ | $u_5$ | *TP53* mutation |

**Table 2:**
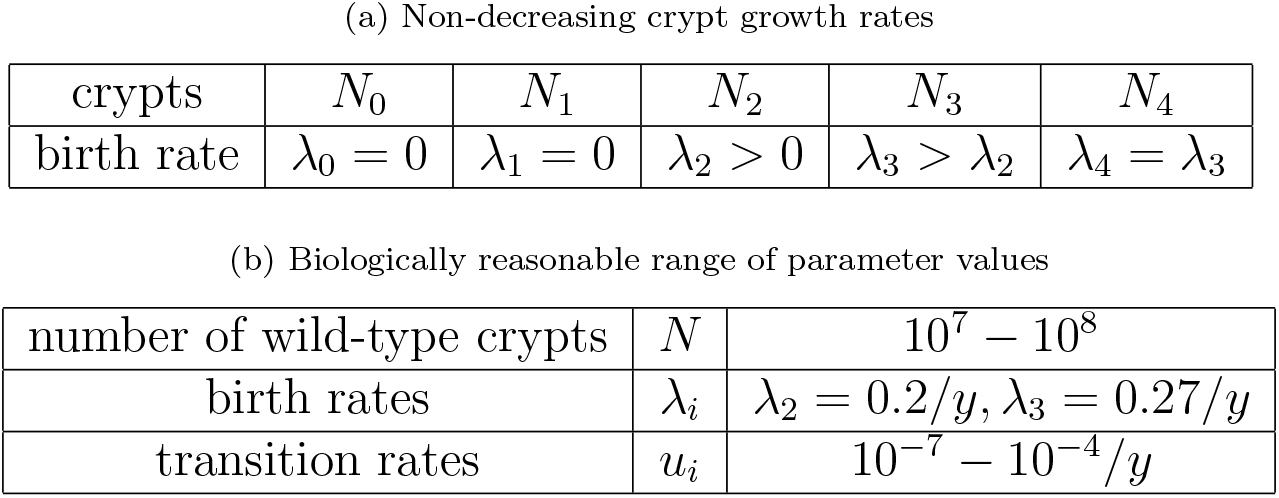
Parameter values for the model of CRC initiation.

|  |  |  |  |  |  |
| --- | --- | --- | --- | --- | --- |
| crypts | $N_0$ | $N_1$ | $N_2$ | $N_3$ | $N_4$ |
| birth rate | $\lambda_0 = 0$ | $\lambda_1 = 0$ | $\lambda_2 > 0$ | $\lambda_3 > \lambda_2$ | $\lambda_4 = \lambda_3$ |

Table 2: Parameter values for the model of CRC initiation.
|  |  |  |
| --- | --- | --- |
| number of wild-type crypts | $N$ | $10^7 - 10^8$ |
| birth rates | $\lambda_i$ | $\lambda_2 = 0.2/y, \lambda_3 = 0.27/y$ |
| transition rates | $u_i$ | $10^{-7} - 10^{-4}/y$ |

This branching process model can be summarized as

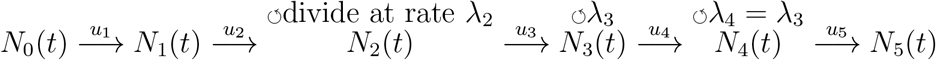

The first generation of type-*i* crypts (all type-*i* crypts produced directly from type-(*i* – 1) crypts) follows an inhomogeneous Poisson process, denoted by *K_i_*(*t*), with instantaneous intensity *u_i_N*_*i*−1_(*t*). The transition times are the founding times of new type-*i* subclones. They are defined as

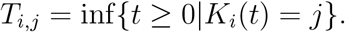

At the *j*-th transition time, the newly formed type-*i* crypt founds a linear birth-death process with birth rate λ_*i*_ and death rate *u*_*i*+1_ (transition rate to the next type). Let *Z*_*i,j*_(*t*) denote this branching process where *t* is measured from the founding time. We also call *Z_i,j_*(*t*) the *j*-th lineage or *j*-th subclone of type-*i* individuals. Let *Z*^(*i*)^ represent a single type-*i* individual. The transition scheme is

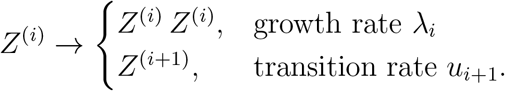

Suppose *Z_i_*(*t*) is a linear birth-death process intiated by a single cell and *Z_i_*(*t*) = 0 for *t* < 0. Then measured from the starting time of CRC, *Z_i,j_*(*t*) is merely *Z_i_*(*t*) shifted by *T_i,j_*. Thus

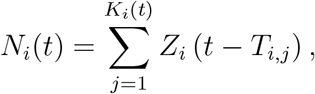

*T_i,j_* ≤ *t*, for 1 ≤ *j* ≤ *K_i_*(*t*). Here we note that since *u*_*i*+1_ is small, we treat all lineages of type-*i* individuals as pure birth processes when type-*i* has a positive birth rate. The system initially consists N wild-type (type-0) crypts, and we seek to estimate the waiting times for the first type-*i* crypt which is defined by

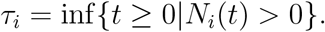

To verify our analytic results, we developed Monte Carlo simulations of a multi-type branching process model based on the Gillespie algorithm [35]. Parameter values for our model come from Paterson et al. [11], and their typical ranges are listed in Table 2.

## 3. General results for a multi-type branching process

In this section, we present and extend a common approach for computing the waiting time distribution of the first type-*i* individual in a multi-type branching process in which type-*i* individuals are produced from type-(*i* – 1) individuals at rate *u_i_*, and each newly formed type-*i* individual starts a new pure birth process with growth rate λ_*i*_. Even though multi-type models may contain many types, mutation from one type to the next can be treated as a two-type branching process. From this perspective, the process containing types 1,…, *n* can be treated as a sequence of *n* – 1 two-type branching processes, with the terminal state from one two-type branching process serving as the initial state for the next. We will show that when large time limits exist for type-*i* populations, it suffices to analyze two-type (initial and terminal type) branching processes (*N_i_*(*t*), *N*_*i*+1_(*t*)), leading to a unified approach. When a large time limit does not exist, we introduce an different approach (see Section 3.4).

In previous studies, the asymptotic population dynamics of two-type branching processes have been analyzed by decomposing the population of the initial type into the product of a deterministic, time-dependent exponential function and a time-invariant random variable [10, 18, 36]. This approximation greatly simplifies the computation of the waiting time distribution by making the stochastic component of the dynamics time-invariant. However, these studies typically consider the case of a wild type with a net growth rate λ_*i*_ strictly greater than zero. In contrast, the wild-type population in our model is not supercritical and in fact (slowly) declines. This, along with allowing λ_*i*_ = λ_*i*+1_, causes non-exponential growth of mutant populations. In this section, we extend previous approaches [10, 18, 36] so that the time-deterministic component can belong to a class of non-exponential functions, which enables us to deal with the non-exponentially growing populations (type-0 and type-1). Furthermore, we review canonical expected value formulas that will be applied in later sections. Similarly to previous approaches [10, 18] we will assume that the mutation rates from one type to the next are small.

### 3.1. Expected value and Laplace transform

Throughout Section 3, we will be concerned with a multi-type branching process (*N*_0_(*t*), *N*_1_(*t*), …)_*t*≥0_, where *N_i_*(*t*) denotes the number of type-*i* individuals at time *t*. The process is started at time zero, with some positive number of type-0 individuals. type-*i* individuals can divide with rate λ_*i*_ ≥ 0 and mutate to type-(*i* + 1) with rate *u_i_* > 0.

We start with a lemma that provides the Laplace transform of *N_i_*(*t*) conditional on the population of its precursor, *N*_*i*−1_(*t*).

#### Lemma 3.1.

*Let Z_i_*(*t*) *be the number of type-i individuals in a pure-birth process that starts with Z_i_*(0) = 1 *individuals at time t* = 0. *Then*

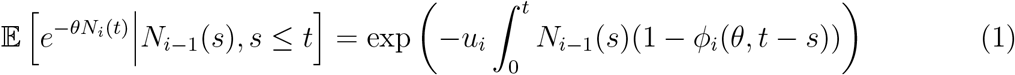

*where* 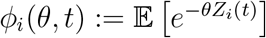.

This is a well-known result for multi-type branching processes. One can prove it by following the procedure of Lemma 2 in [10], replacing the start time with *s* = 0.

The expected value formula for the number of type-*i* individuals can be derived from Lemma 3.1.

#### Theorem 3.2.

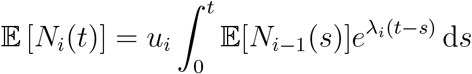

Taking the derivative of equation (1) and evaluating at *θ* = 0 gives the relationship between expected values.

### 3.2. Large time limits

#### Lemma 3.3.

*Consider* (*N*_*i*−1_(*t*), *N*,(*t*))_*t*≥0_. *Suppose N_i_*(*t*) *is the population of type-i crypts in the multi-type branching system where the population of type*-(*i* – 1) *crypts is given by N*_*i*−1_(*t*) ≥ 0, *then*

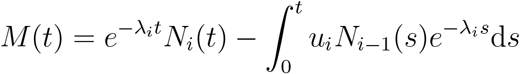

*is a martingale. If*

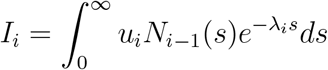

*has a finite expectation, then*

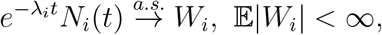

*as t* → ∞. *Additionally, if e*^-λ_*i*_*t*^*N_i_*(*t*) *is uniform integrable, then*

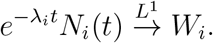

*This implies*

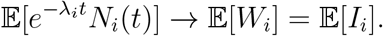

If the first condition from the statement of the Lemma holds (i.e. if *I_i_* has finite expectation) Lemma 3.3 provides a method of obtaining the long-term behavior of *N_i_* using the limiting random variable *W_i_*. In that case, we have *e*^-λ_*i*_*t*^*N_i_*(*t*) → *W_i_*, and for large time *t*, *e*^λ_*i*_*t*^*W_i_* should be a good approximation of the stochastic process *N_i_*(*t*). The importance of *N_i_*(*t*) ≈ *e*^λ_*i*_*t*^*W_i_* is that it separates a stochastic process into a deterministic function *e*^λ_*i*_*t*^ and a time-independent random variable *W_i_*.

If, in addition, the second condition (*e*^-λ_*i*_*t*^ *N_i_*(*t*) is uniform integrable) holds, then the expected value of the limiting random variable *W_i_* is obtainable. In that case, we have 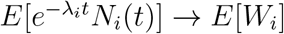, which makes the large time approximation *N_i_* (*t*) ≈ *e*^λ_*i*_*t*^*W_i_* reasonable in terms of the first moment.

### 3.3. Estimating waiting times using large time limits

Let *τ_i_*, (1 ≤ *i* ≤ *n*) be the waiting time of the first type-*i* individual in a multi-type branching process. At time *s* ≥ 0, the arrival rate of type-*i* individuals is *u_i_N*_*i*−1_(*s*). Conditional on the trajectory of *N*_*i*−1_(*s*) for 0 ≤ *s* ≤ *t*, the probability of *τ_i_* can be written as

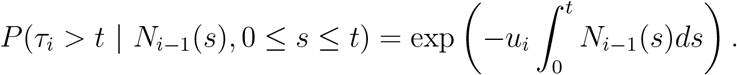

The functional form of *N*_*i*−1_(*s*) is generally unknown and potentially complicated. One way of evaluating this integral is to approximate *N*_*i*−1_(*s*) by the product of a deterministic time-dependent growth and a time independent random variable. For example, let *N*_0_(*t*) = *Z*_0_(*t*), a pure birth two-type branching process that starts with a single individual. It is well-known that *e*^-λ_0_*t*^*Z*_0_(*t*) → *W*_0_ ∼ Exponential(1) [10]. A classical approximation is *N*_0_(*s*) ≈ *e*^λ_0_*t*^*W*_0_ where the deterministic time-dependent growth is characterized by *e*^λ_0_*t*^ and time independent random variable is *W*_0_. Applying this approximation yields

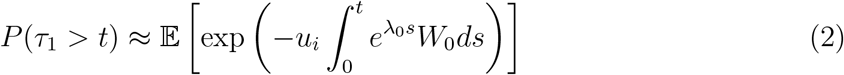

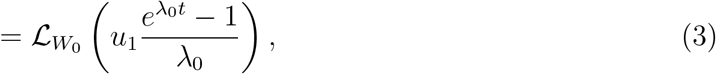

vhere 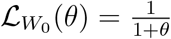 is the Laplace transform of *W*_0_. Let *f*_0_(*t*) = *e*^λ_0_*t*^ be the time determinstic function. Then the above approximation also holds if a sub-exponential term is added to *f*_0_. In other words, we have many reasonable options for *f*_0_. In later sections, we consider two specific deterministic functions,

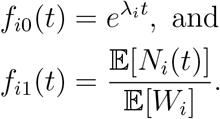

In the example mentioned above, we have

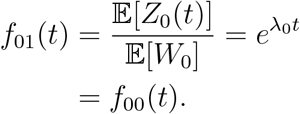

However, in our model when *i* > 1, *f*_*i*0_ ≠ *f*_*i*1_. Later we will see an approximation with *f*_*i*1_ is more precise than that with *f*_*i*0_.

The second observation from this example is that the approximation (2) relies on the Laplace transform of *W*_0_. However, one cannot obtain a closed form for the Laplace transform of *W_i_,i* ≥ 2. In this case, an alternative approach is to approximate *W_i_* by another random variable *V*_i_ that has a a closed form Laplace transform. Such *V_i_* can be found by approximating *N*_*i*−1_, as described in Section 4.

#### Proposition 3.4.

*Let* (*N*_*i*−1_ (*t*), *N_i_*(*t*)) *be a two-type process such that N_i_*(0) = 0 *and type-i individuals are produced by type*-(*i* – 1) *individuals with rate u_i_N*_*i*−1_(*t*),*u_i_* > 0. *Suppose that* 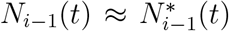 *such that* 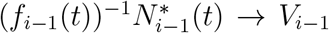 *as t* → ∞. *Then the waiting time distribution of the first type-i individual can be approximated by*

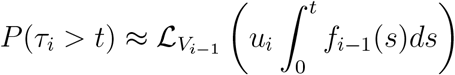

*where* 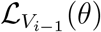 *is the Laplace transform of random variable V*_*i*−1_.

Applying the approximations gives us

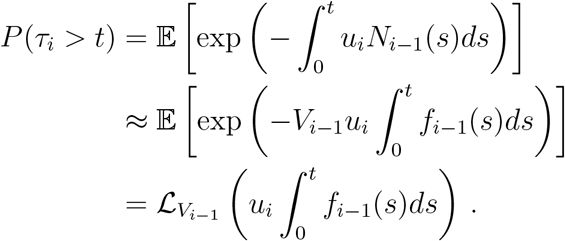

From Proposition 3.4, we see that the waiting time (*τ_i_*) distribution depends on the Laplace transform of *V*_*i*−1_. Here we present an iterative method for computing the Laplace transforms of *V_i_*.

#### Lemma 3.5.

*Let* 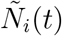 *be type-i individuals in the system with* 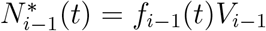. *Suppose* 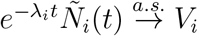, *then*

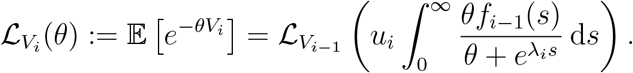

### 3.4. An inhomogeneous Poisson process approximation

Proposition 3.4 estimates the arrival time of type-*i* individuals by the large time limit of the previous type (*V*_*i*−1_). However, there are situations when the random variable *V*_*i*−1_ does not exist or when *f*_*i*−1_ *V*_*i*−1_ fails to provide a precise first arrival time of *N_i_*. More generally, we may only have a good large time limit *V_j_* for some type-*j* where *j* < *i*, so that we do not have reliable limits for type-(*j* + 1) through type-(*i* – 1). To deal with this situation, we introduce a method that uses the large time limit of each independent lineage to ‘skip’ *V*_*j*+1_ through *V*_*i*−1_. First, note that given *V_j_*, we can approximate the birth rate of type-(*j* + 1) lineages, using the well-known fact that for each type-(*j* + 1) lineage,

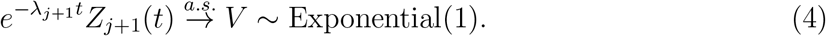

Next, we want to find the likelihood of each type-(*j* + 1) lineage producing at least a single type-*i* individual. Recall that a type-(*j* + 1) lineage is a simple birth process initiated by a single type-(*j* + 1) individual that grows at rate λ_*j*+1_. Suppose we have a type-(*j* + 1) lineage that was started by a single type-(*j* + 1) individual at time *s*. We define 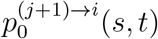 to be the probability that no type-*i* individual is produced by time *t* by this type-(*j* + 1) lineage.

At time *s*, type-(*j* + 1) crypts are producing at rate 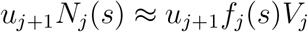. Each type-(*j* + 1) lineage (present at time *s*) has a probability 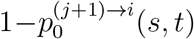 to produce at least a single type-*i* crypt (at time *t*). Finally, for fixed *t*, we approximate the process of producing type-*i* individuals as an inhomogeneous Possion process with rate 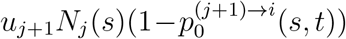. This implies

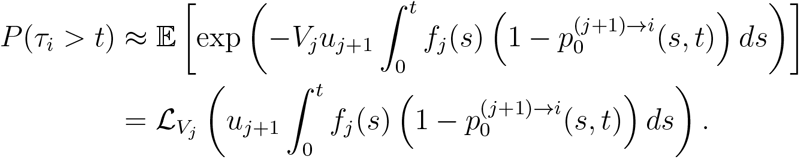

This approach is summarized in the following proposition.

#### Proposition 3.6.

*Let* (*N_j_* (*t*), *N*_*j*+1_(*t*), ⋯, *N_I_*(*t*)) *be a* (*i* – *j* + 1)-*type process such that N_k_*(0) = 0, *j* < *k* ≤ *i and type-k individuals are produced by type*-(*k* – 1) *individuals with rate u_k_N*_*k*-1_(*t*),*u_k_* > 0. *Suppose that* 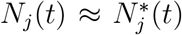 *such that* 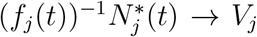. *Then the waiting time distribution of type-i crypts can be approximated by*

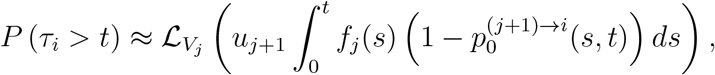

*where* 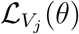 *is the Laplace transform of V_j_*.

Proposition 3.4 is consistent with Proposition 3.6, and one can treat Proposition 3.4 as a single-step version of Proposition 3.6.

To compute 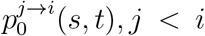, we use the iterative relationship between 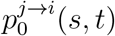 and 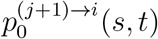. This is provided by the following proposition.

#### Lemma 3.7.

*For i* < *j*,

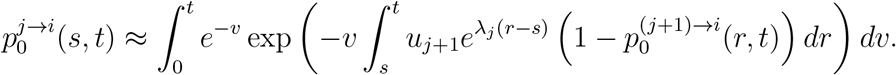

*with* 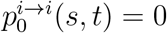.

## 4. Population dynamics of type-*i* crypts

In this section we derive the asymptotic growth dynamics of the clonal subpopulations (or types) along the path to CRC initiation. For each type *i*, the population size *N_i_*(*t*) is a counting process indexed by *t* > 0. The first two types, *N*_0_(*t*) and *N*_1_(*t*), are well approximated by time-deterministic functions. Starting with these, we sequentially derive the large-time population dynamics for the later types, leveraging results for two-type branching processes, (*N_i_,N*_*i*+1_), 0 ≤ *i* ≤ 4. The success of this approach relies on our ability to approximate the large-time growth dynamics by the product of a time-deterministic function and a time-invariant random variable. The existence of such a decomposition depends on the martingale properties of the counting processes *N_i_*(*t*).

### 4.1. Type-0

Wild-type human crypts (i.e. type-0 crypts) rarely divide [32], hence we set the growth rate of the wild-type crypts to zero (λ_0_ = 0). These wild-type crypts transition into type-1 crypts at rate *u*_1_*N*_0_(*t*) after losing one copy of the *APC* gene.

#### Proposition 4.1.

*N*_0_(*t*) *is a pure death process with death rate u*_1_ *and initial condition N*_0_(0) = *N. Its generating function at time t is*

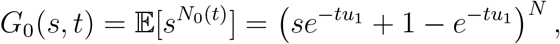

*and the expectation and variance are*

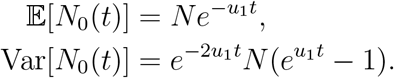

For a detailed description of a single-type birth-death process, one can refer to chapter 3, section 5 in [37]. Typical biologically realistic parameter values are *u*_1_*t* ~ 10^-2^ ≪ 1 and *N* ~ 10^8^, which leads to

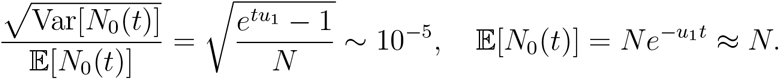

Motivated by this result, we approximate the population of the wild-type crypts by its expectation

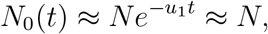

in which the second approximation is justified because *u*_1_*t* ≪ 1. Therefore, the waiting time of *N*_1_(*t*), *τ*_1_ ~ Exponential(*u*_1_ *N*).

### 4.2. Type-1

A type-1 crypt is produced when a healthy crypt (type-0) loses one copy of the *APC* gene. This mutation does not lead to clonal expansion [33], so the growth rate of type-1 crypts, λ_1_ = 0. While being produced by wild-type crypts, type-1 crypts mutate into type-2 at rate *u*_2_*N*_1_(*t*), losing both copies of the *APC* gene. We assume that the population outflows are negligible. Initially, there are no type-1 crypts, i.e., *N*_1_(0) = 0.

#### Proposition 4.2.

*N*_1_(*t*) = *N* – *N*_0_(*t*) *has the following expectation and variance*:

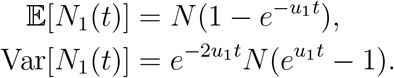

This can be derived by observing *N*_0_(*t*) + *N*_1_(*t*) = *N*, since the outflows from type-1 to type-2 are negligible. Therefore 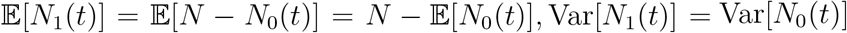.

To approximate *N*_1_(*t*), we consider the case when the wild-type population is effectively constant. In this case, *N*_1_(*t*) can be treated as a Poisson process with intensity *u*_1_*N*. According to the properties of a Poisson process, the approximated expectation and variance are

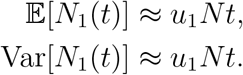

Notice that the approximated expected value and variance are consistent with the true expected value and variance. Since *u*_1_*t* ~ 10^-2^ ≪ 1 is small, we have

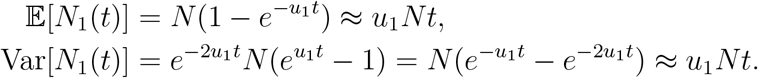

Additionally, observe that

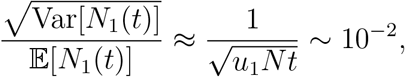

and so we approximate *N*_1_(*t*) by a deterministic function

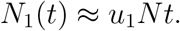

Thus the waiting time distribution of type-2 crypts is simply estimated as

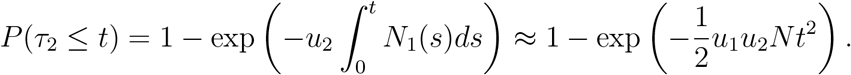

### 4.3. Type-2

After losing both copies of the *APC* genes, type-1 crypts mutate into type-2 crypts. The *APC* inactivation provides a fitness advantage to type-2 crypts [33]. Thus for type-2, we have a positive growth rate λ_2_ > 0. Meanwhile, the *KRAS* oncogenes are activated at rate *u*_3_*N*_2_(*t*), mutating type-2 crypts into type-3 crypts. These outflows are ignored when considering the population of *N*_2_(*t*). At time *t* = 0, there are no type-2 crypts, i.e *N*_2_(0) = 0. We begin with the expectation of *N*_2_(*t*). Using Theorem 3.2, we compute

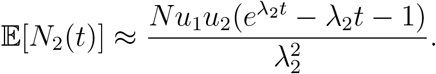

The approximation is obtained by noting that *u*_1_ ~ 10^-4^ ≪ λ_2_ ~ 10^-1^ and *u*_1_*t* ~ 10^-2^ ≪ 1. The following large-time asymptotic limit exists for *N*_2_(*t*).

#### Theorem 4.3.

*e*^-λ_2_*t*^*N*_2_(*t*) → *W*_2_ *a.s. and in L*^1^ *with*

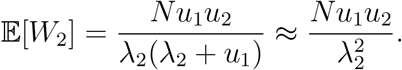

To characterize the distribution of the limiting random variable, we consider 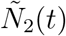, the population of type-2 crypts produced by 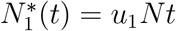.

#### Theorem 4.4.

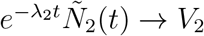 *a.s. and in L*^1^ *with*

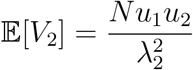

*and*

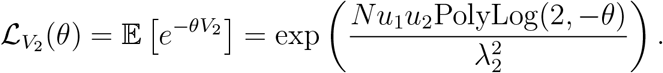

#### Corollary 4.5.

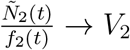 *a.s. and in L*^1^ *for all*

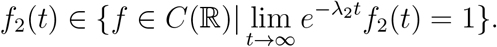

### 4.4. Type-3

Type-3 crypts are produced by type-2 crypts through activation of the *KRAS* oncogene, which increases the fission rate of mutated cells [34]. Thus the birth rate has a positive increment, i.e. λ_3_ > λ_2_. At a small rate *u*_4_*N*_3_(*t*), type-3 crypts lose one copy of the TSG *TP53* and mutate into type-4 crypts. The initial population is *N*_3_(0) = 0. Using Theorem 3.2, the expected value of *N*_3_(*t*) is

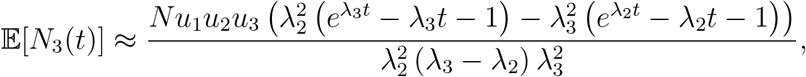

where the approximation is made by observing λ_*i*_ + *u*_1_ ≈ λ_*i*_ and *u*_1_*t* ~ 10^-2^.

#### Theorem 4.6.

*e*^-λ_3_*t*^*N*_3_(*t*) → *W*_3_ *a.s. and in L*^1^ *with*

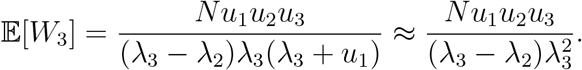

Consider a system with 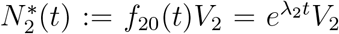 and let 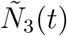 denote the number of type-3 crypts in this system. The following large-time limit holds.

#### Theorem 4.7.

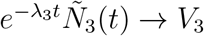 *a.s. and in L*^1^ *with*

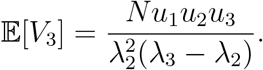

For other potential *f*_2,*k*_’s, the Laplace transform changes accordingly.

#### Corollary 4.8.

*Let* 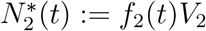 *where*

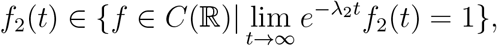

*and let* 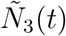 *denote type-3 crypts in this system. Then* 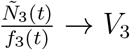 *a.s. and in L*^1^ *with*

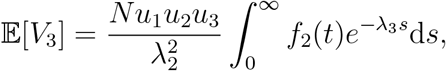

*and*

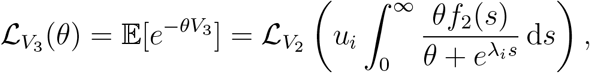

*for all*

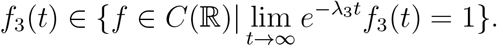

Note that 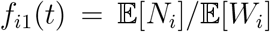 is more precise than *f*_*i*0_(*t*) = *e*^λ_*i*_*t*^. By Corollary 4.8, 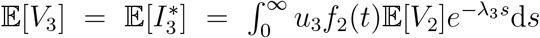, depends on *f*_2_(*t*) and 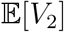. Since 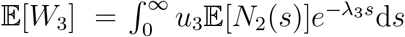 and 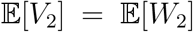, *setting* 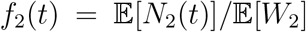 equates 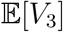 and 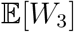. Indeed, when using *f*_2_(*t*) = *f*_20_(*t*) = *e*^λ_2_*t*^, the resulting random variable *V*_3_ has an expectation different from *W*_3_.

### 4.5. Type-4

Since only one copy of the *TP53* gene has been lost, type-4 does not confer a growth advantage [30]. Thus the division rate of type-4 crypts λ_4_ = λ_3_. The dynamics of type-4 crypts also include negligible outflows at rate *u*_5_*N*_4_(*t*). And the starting condition is *N*_4_(0) = 0. In this case, we note that using *e*^-λ_3_*t*^ to scale the growth process fails to provide a large-time limiting random variable. Even though

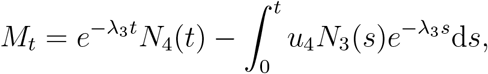

is still a martingale, Lemma 3.3 does not apply. We have seen that 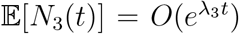 as *t* → ∞, and so

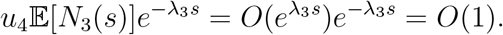

This implies the first condition in Lemma 3.3 is invalid, that is as *t* → ∞

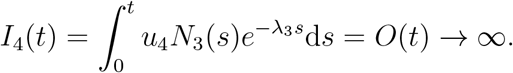

Therefore we cannot derive a large time limit of type-4 individuals. Instead, we will estimate the waiting time of type-5 crypts by the inhomogeneous Poisson approximation discussed in section 3.4. To this end, we compute 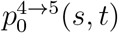 and 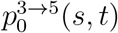, corresponding to estimating *τ*_5_ by *V*_3_ and *V*_2_ respectively,

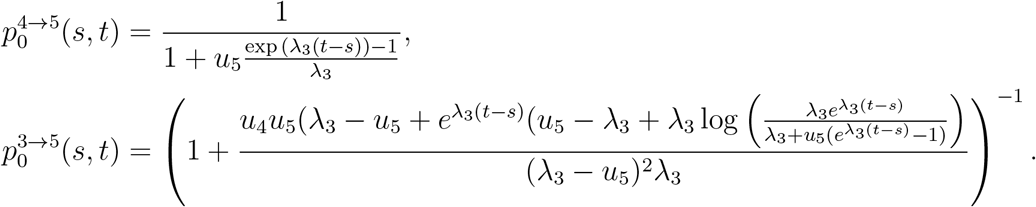

## 5. Estimating waiting times for colorectal cancer initiation

In this section, we compare results obtained from exact computer simulations of the multitype process with our analytic results. We define *p_i_*(*t*) as our estimate of the cumulative distribution function of *τ_i_*, the waiting time to the first type-*i* crypt. For all figures in this section, parameter values are given in Table 3, and follow estimates from Paterson et al [11].

**Table 3:** Estimates of parameter values for colorectal cancer initiation from Paterson et al. [11].

| Crypt $N_i$ | $N_0$ | $N_1$ | $N_2$ | $N_3$ | $N_4$ |
| --- | --- | --- | --- | --- | --- |
| Initial population (crypts) | $N = 1 \times 10^8$ | 0 | 0 | 0 | 0 |
| Birth rate $\lambda_i$ (crypts/year) | 0 | 0 | 0.2 | 0.27 | 0.27 |
| Transition rate $u_{i+1}$ (crypts/year) | $2.86 \times 10^{-4}$ | $1.06 \times 10^{-5}$ | $9.00 \times 10^{-7}$ | $1.36 \times 10^{-4}$ | $4.56 \times 10^{-7}$ |

### 5.1. Waiting times to type-1 and type-2

For *τ*_1_ and *τ*_2_, the analytic results are listed in the previous section. Recall that the population of wild-type (type-0) crypts are estimated to have a fixed population *N*. Thus,

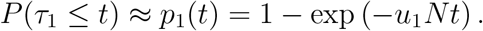

For *τ*_2_, recall that we approximate the population of type-1 crypts by a time deterministic function *u*_1_*Nt*.

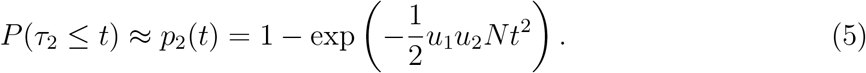

### 5.2. Waiting time to type-3

To compute the distribution functions of *τ*_3_, *τ*_4_ and *τ*_5_, we need to apply the approximation in Proposition 3.4. Hence, we want to find *f*_2_, *f*_3_. Recall that we consider two functions for each subpopulation:

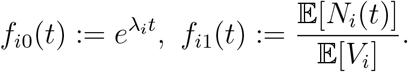

The approximation for the distribution *τ*_3_ is

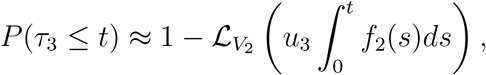

where 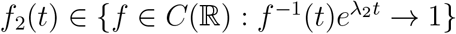. First, consider *f*_20_(*t*) = *e*^λ_2_*t*^. The corresponding distribution function is

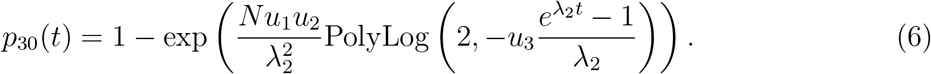

In the second case,

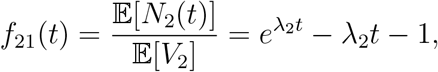

and the corresponding distribution is

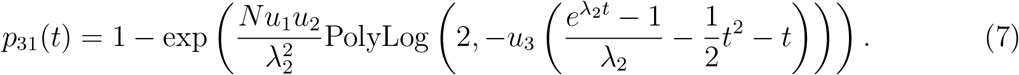

By design, the first moment of 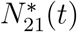 is identical to that of *N*_2_(*t*),

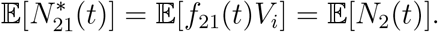

Both (6) and (7) agree with the simulated distributions for *t* > 40 (See Fig 2). However, in the intermediate regime where t is small, one can observe that *p*_31_ is more accurate than *p*_30_.

**Figure 1:**
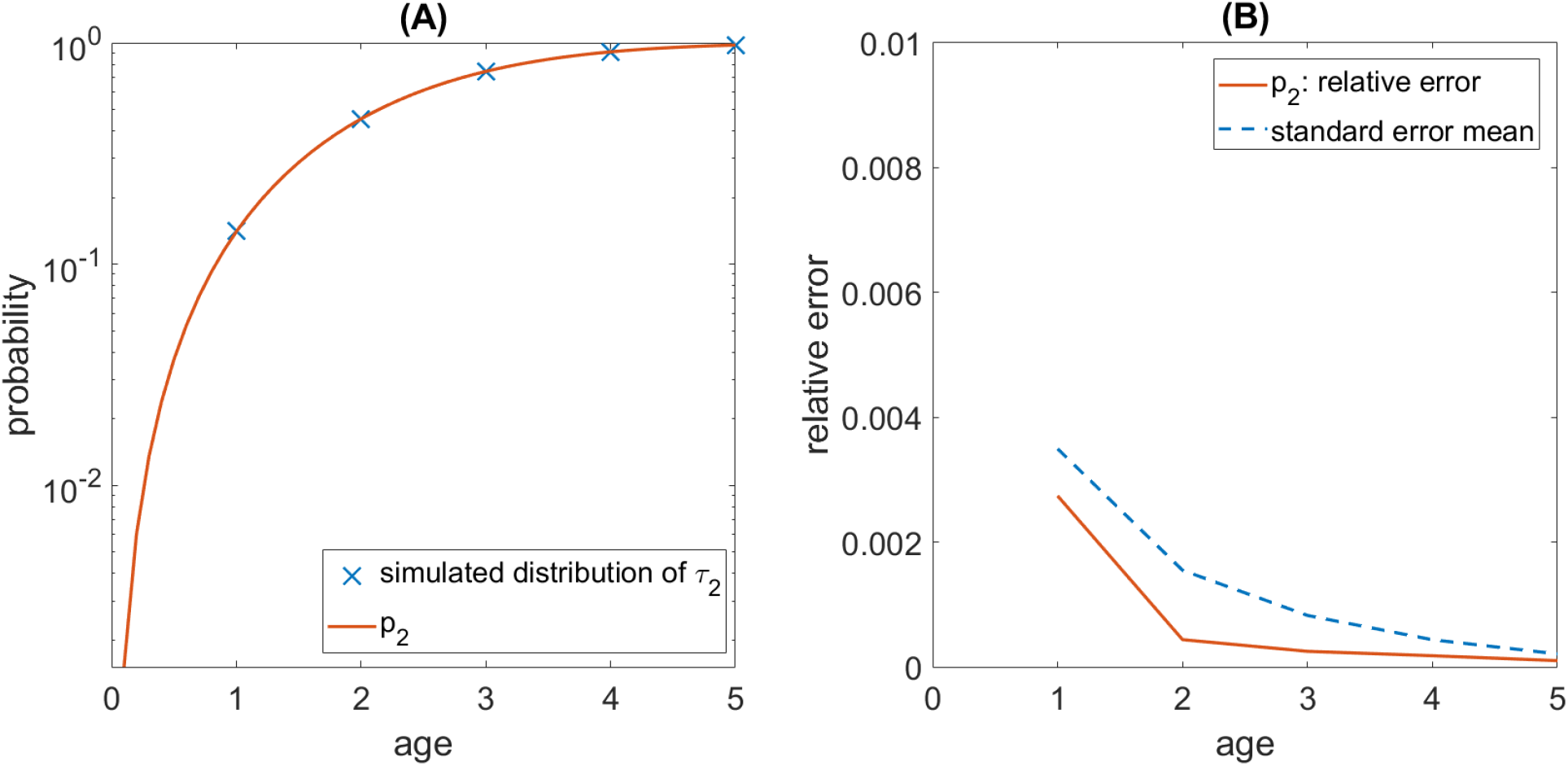
(A) Comparison of the analytic cumulative distribution function of *τ*_2_, the waiting time to the first type-2 crypt (*p*_2_, equation (5)), and the simulated distribution of *τ*_2_ across 5 × 10^5^ runs. (B) Dashed line shows the standard error of the mean obtained from simulations. Solid line is the relative error of the analytic result.

**Figure 2:**
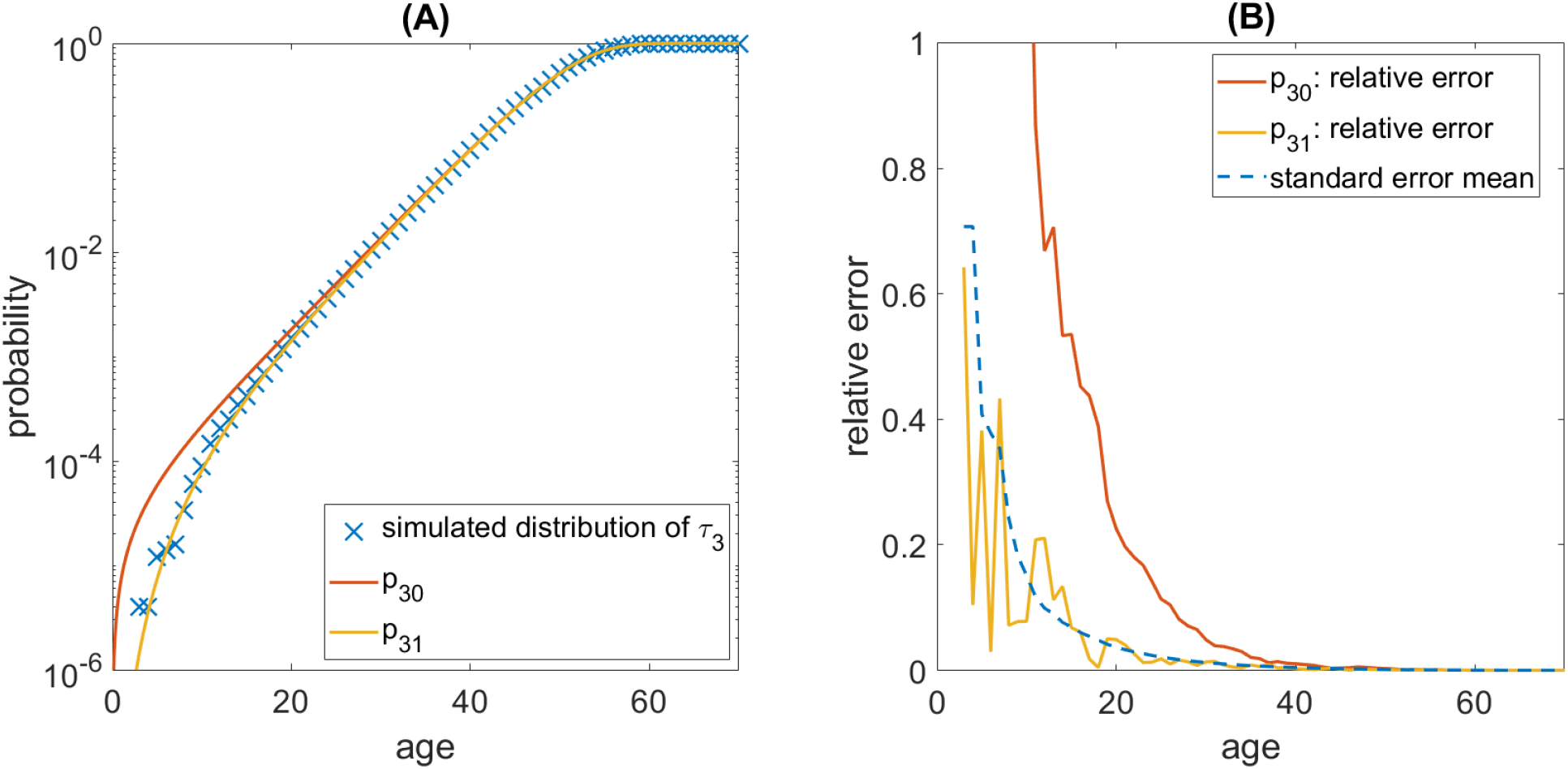
(A) Comparison of the analytic cumulative distribution functions of *τ*_3_, the waiting time to the first type-3 crypt, and the distribution of *τ*_3_ across 5 × 10^5^ simulation runs. *p*_30_ and *p*_31_ are distribution functions obtained from equations (6) and (7) respectively. (B) Dashed line shows the standard error of the mean of the simulation. Solid lines are the relative errors of the analytic results.

### 5.3. Waiting time to type-4

Considering Proposition 3.6, we have several options to compute *τ*_4_, the waiting time to the first type-4 crypt. Specifically, we may compute the distribution of *τ*_4_ in two ways: (i) compute the distribution directly using 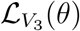; and (ii) skip *V*_3_ and compute the distribution using 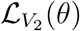 and 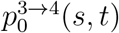. In these approaches, by Corollary 4.8, the time deterministic growth functions *f*_2_, *f*_3_ can be any functions that have large time limits identical to the corresponding exponential functions, *e*^λ_2_*t*^ and *e*^λ_3_*t*^. Among different choices of the growth functions, we typically consider two specific cases, *f_i_* = *f*_*i*0_ = *e*^λ_*i*_*t*^ and 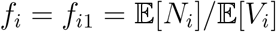.

If *V*_3_ is not skipped, 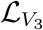 depends on the choice of *f*_2_. Since *f*_21_ leads to more accurate computations of *τ*_2_, we use *f*_21_ to compute 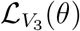, that is

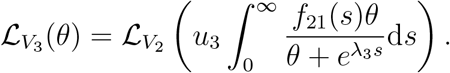

For *f*_3_, we consider two candidate functions,

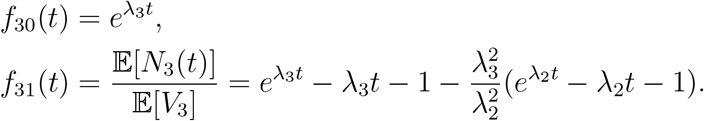

The first, *f*_30_(*t*), has a simpler form corresponding to the leading order of 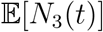, with

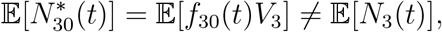

and the second, *f*_31_(*t*), matches the expected value of 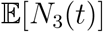

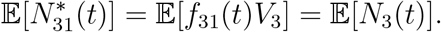

Next, we define

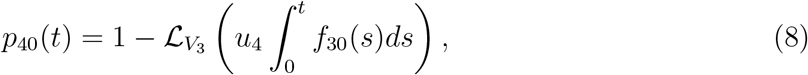

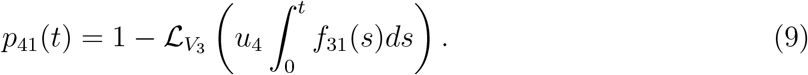

On the other hand, if we skip *V*_3_ and use the inhomogeneous approximation in section 3.4, we can consider two types of *f*_2_’s:

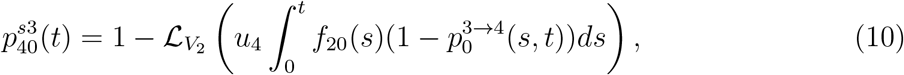

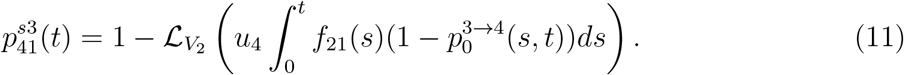

We note that 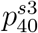 and 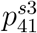 have closed form expressions, which are omitted here and listed in the Appendix. The results are shown in Fig 3. We observe that skipping *V*_3_ improves the accuracy of the results for large time *t*, while including more terms in *f_i_* generally improves the accuracy of the approximation.

**Figure 3:**
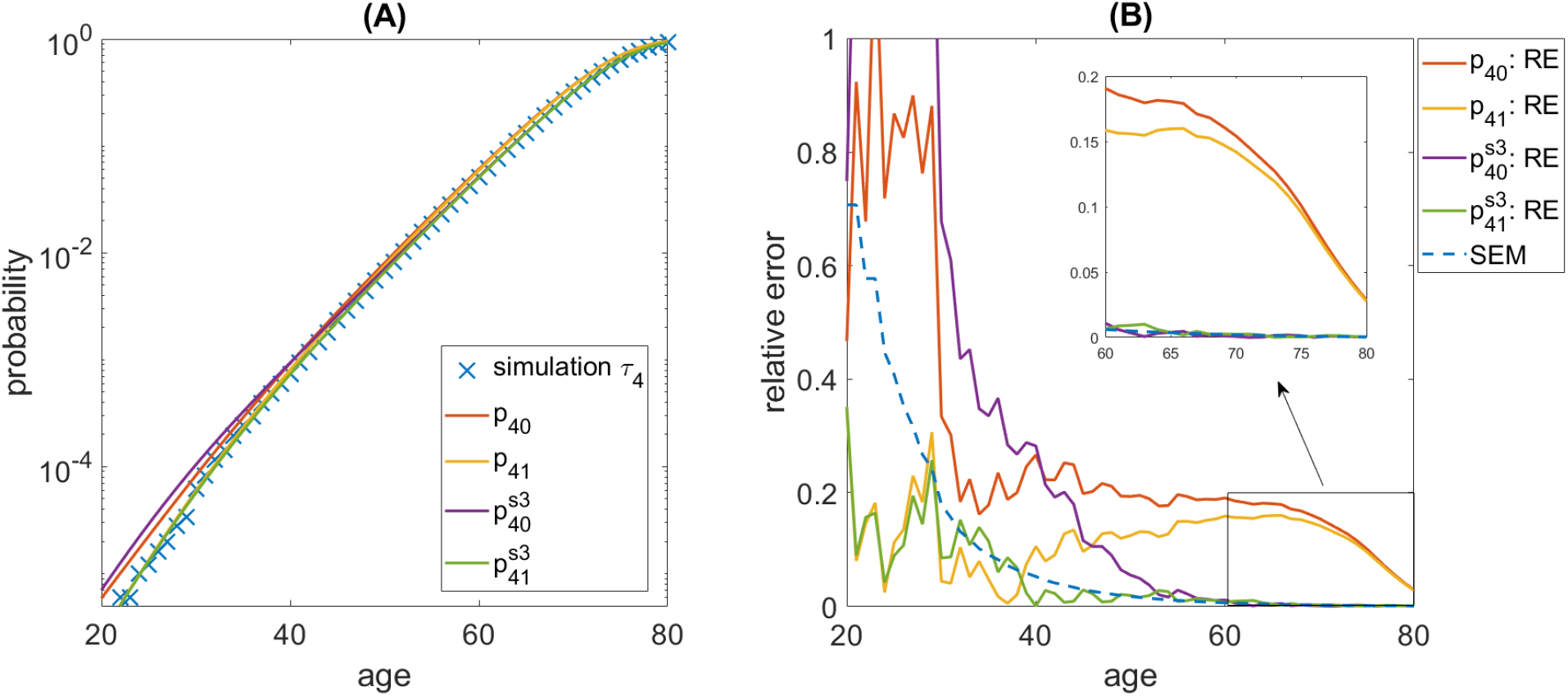
(A) Comparison of analytic cumulative distribution functions of *τ*_4_, the waiting time to the first type-4 crypt, and the distribution of *τ*_4_ across 5 × 10^5^ simulation runs. *p*_40_ (equation (8)) and *p*_41_ (equation (9)) are analytic distribution functions derived from the Laplace transform *V*_3_. 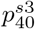 (equation (10)) and 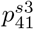 (equation (11)) are analytic distribution functions derived by skipping *V*_3_ and using the Laplace transform of *V*_2_. (B) Dashed line shows the standard error of the mean of the simulation. Solid lines are the relative errors of the analytic results.

### 5.4. Waiting time to type-5

For *τ*_5_, the waiting time to the first type-5 crypt, we consider two approaches: (i) compute the distribution using the large-time limit of type-3 crypts, 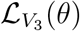, and 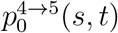 (effectively skipping the large-time limit of type-4 crypts); and (ii) skip large-time limits of both type-4 and type-3 crypts, and compute the distribution using 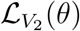 and 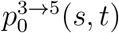. Note that the time-deterministic growth functions *f*_2_ and *f*_3_ which appear in the long-time limits of type-2 and type-3 crypts can vary across these approaches (see Corollary 4.8). However, the most accurate results are obtained when 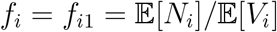.

The corresponding approximations of the distribution of *τ*_5_ resulting from approaches (i) and (ii) are

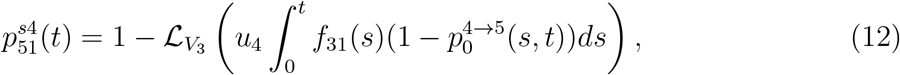

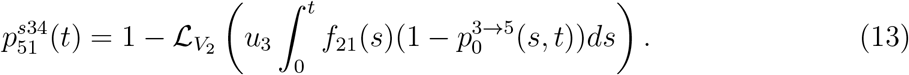

The first result, 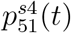, has an explicit form given by equation (B.1), while the second result, 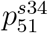 (given by equation (B.3)), to the best of our knowledge, is not an elementary function or a standard special function. Comparison of these two formulas with the waiting time for the first type-5 crypt obtained from exact computer simulations is shown in Fig 4. We observe that 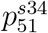 achieves higher accuracy compared to 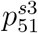, especially at later times (above age 70). In other words, compared with the result incorporating *V*_3_, skipping this stage gives more accurate results. The intuition behind this is that approximating each subclone by its large time limit is more accurate than approximating the total population by its overall large time limit.

**Figure 4:**
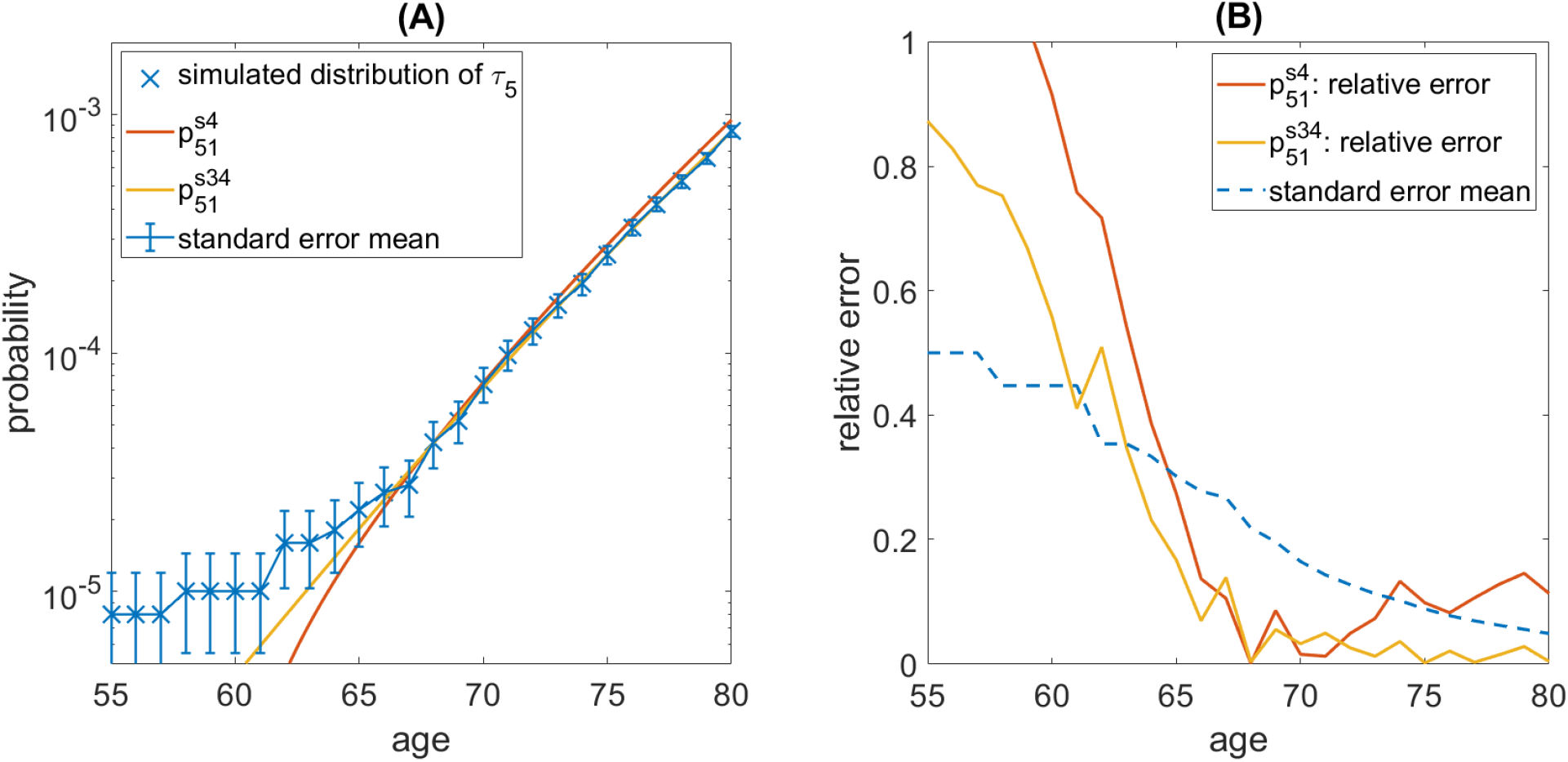
(A) Comparison of the analytic cumulative distribution functions of *τ*_5_, the waiting time to the first type-5 crypt, and the distribution of *τ*_5_ across 5 × 10^5^ simulation runs. 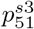 (equation (12)) is derived by skipping type-4 and using the Laplace transform of *V*_3_. 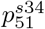 (equation (13)) is derived by using the Laplace transform of *V*_2_ and skipping the Laplace transforms of type-4 and type-3. The error bars represent the standard error of the mean of the simulation. (B) Dashed line shows the standard error of the mean of the simulation. Solid lines are the relative errors of the analytic results.

## Acknowledgements

This work is supported by the National Science Foundation grant DMS-2045166.

## Appendix

## A. Proofs

### Proof of Lemma 3.3

*Proof.* The proof follows that of Theorem 2 in [10]. The only difference is that we have derived a new bound on the martingale. By Lemma 1 in [10],

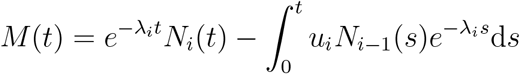

is a martingale. If *I_i_* has a finite expectation, then by the martingale convergence theorem (Theorem 4.2.11 in [38]), the submartingale *X*(*t*) = – *M*(*t*) converges a.s. to some integrable limit *X* as *t* → ∞. Since

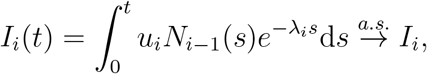

we also have

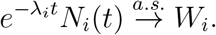

The martingale starts at zero (i.e. *M*(0) = 0), which implies

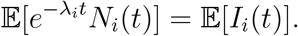

Suppose *e*^-λ_*i*_*t*^*N_i_*(*t*) is uniform integrable, we have (Theorem 4.6.3 in [38])

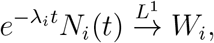

which guarantees

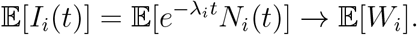

Thus, we have

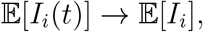

and

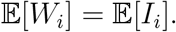

### Proof of Lemma 3.5

*Proof.* To get the Laplace transform of *V_i_*, we start from the star process approximation

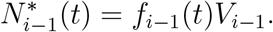

Applying Lemma 3.1 to the 2-type process 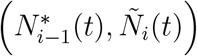 yields

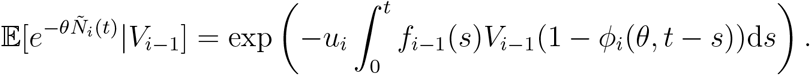

Replacing *θ* with *θe*^-λ_*i*_*t*^,

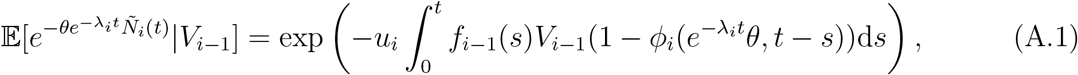

then for each subclone of *N_i_*, by equation (4), we have

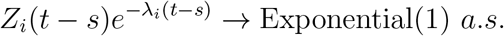

Thus, it follows

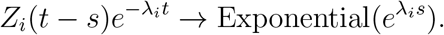

Now consider the following limit involving terms on the right hand side of (A.1),

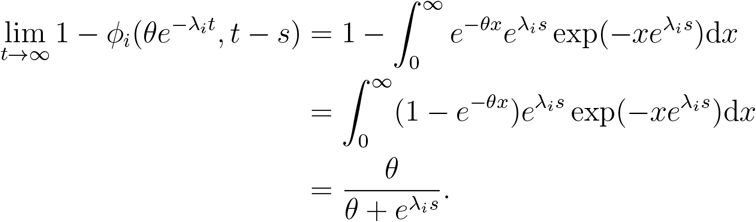

And note that as *t* → ∞, the left hand side of (A.1) yields

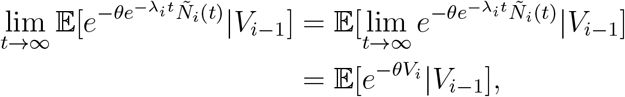

in which switching the limit and the integration is allowed by the dominated convergence theorem. Thus, we can write

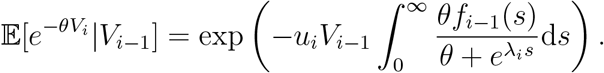

Taking expectation on both sides gives us

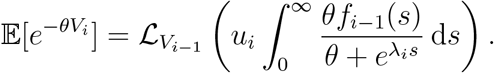

### Proof of Lemma 3.7

*Proof.* For a type-*j* lineage that appears at time *s*, we approximate it by its large time limit. At time *r* > *s*, the population of this lineage would be *Z_j_*(*r*) ≈ *e*^λ_*j*_(*r−s*)^*V*, where *V* ~ Exponential(1). Thus, type-(*j* + 1) individuals that mutated from this lineage are produced at rate

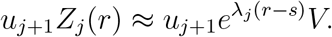

The probability for a type-(*j* + 1) individual to produce at least a type-*i* individual is 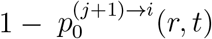. Thus, conditional on *V*, the expected number of type-*i* individuals that were produced by a type-*j* lineage that appeared at time *s* is

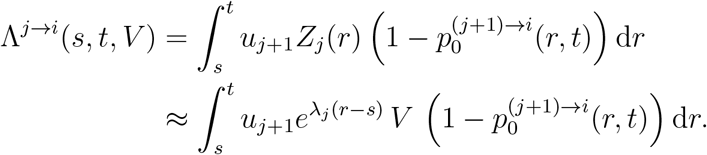

Let *X*^*j*→*i*^(*s, t*) be the number of type-*i* individuals that are produced by a type-*j* subclone appeared at *s*. In the time period [*s,t*], *X*^*j*→*i*^(·,*t*) follows an inhomogeneous Possion process with mean Λ(·,*t, V*). Thus the probability that no type-*i* crypt is produced from this particular type-(*j* + 1) crypt by time *t* is

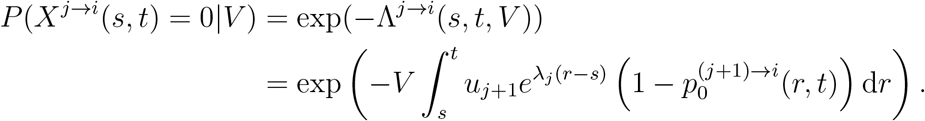

This implies

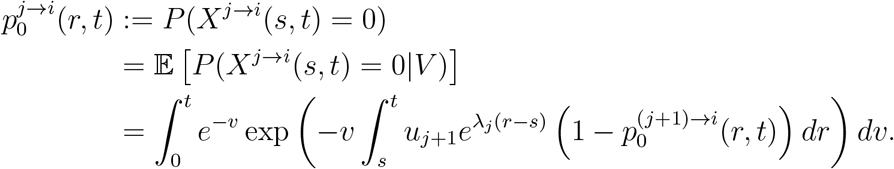

Finally, for *i* = *j*, notice that the founding individual of a type-*j* lineage is of type-*j*. This guarantees that at any time *t* greater than the founding time *s*, the probability of having at least one type-*j* individual is 1. Thus 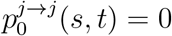.

### Proof of Theorem 4.3

*Proof.* By Lemma 3.3, consider

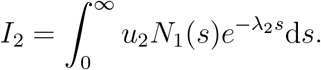

Since

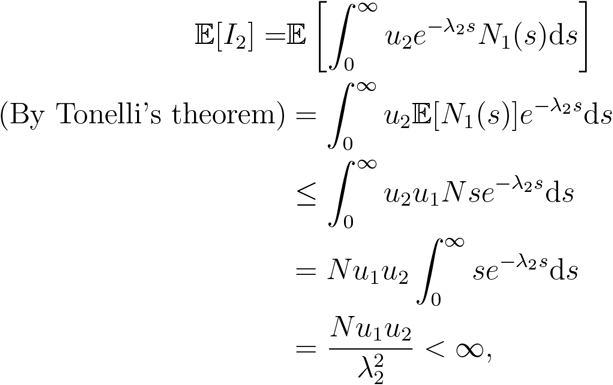

there exists *W*_2_ s.t.

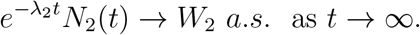

Next, we show uniform integrability so that the 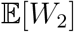 is well-defined. We prove this for *N*_1_, *N*_2_ and *N*_3_ in Lemma A.1. Since *L*^1^ convergence is guaranteed, 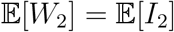, so that

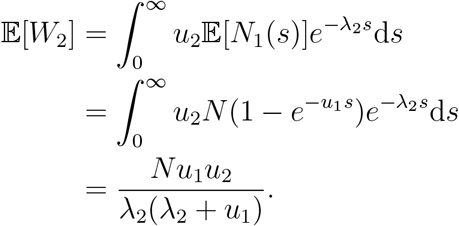

### Proof of Theorom 4.4

*Proof.* Note that 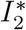 is deterministic and has a finite expected value

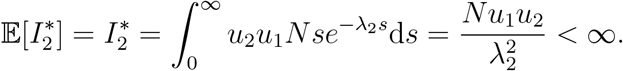

By Lemma 3.3, we must have 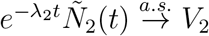. Next, we show uniform integrability which guarantees *L*^1^ convergence. It is shown in Lemma A.2 that all tilde processes in this paper are uniform integrable. This implies 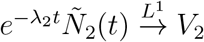 and 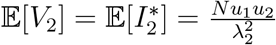. Finally, to compute the Laplace transform of *V*_2_, we plug *f*_2_(*t*) = *e*^λ_2_*t*^ into the formula in 3.5.

### Proof of Corollary 4.5

*Proof.* The convergence directly follows Theorem 4.4 by the fact that

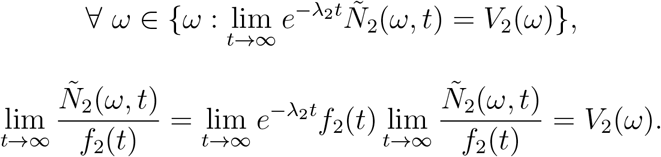

Observe that for *t* > 0 sufficiently large, we have *f*_2_(*t*) > 0 and *e*^λ_2_*t*^/*f*_2_(*t*) < *M*. Note that

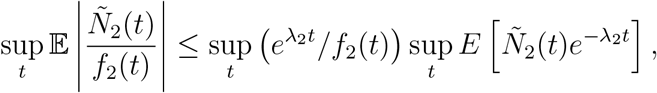

and by Lemma A.2 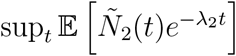 is bounded. Hence 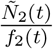 is uniform integrable and the convergence is in *L*^1^.

### Proof of Theorem 4.6

*Proof.* By Lemma 3.3 and Lemma A.1, we need to verify that *I*_3_ has finite expectation.

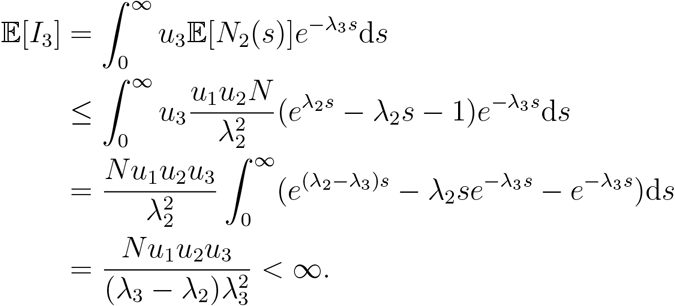

And the expected value of *W*_3_ is given by the expected value of *I*_3_,

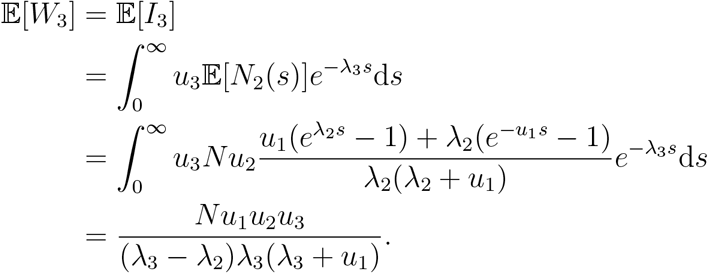

### Proof of Theorem 4.7

*Proof.* By Lemma 3.3 and Lemma A.2, we compute

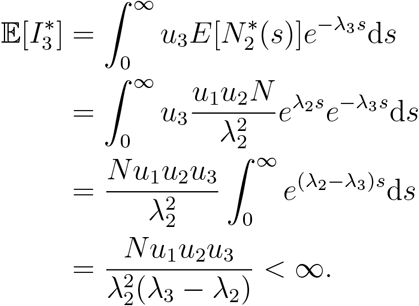

### Proof of Corollary 4.8

*Proof.* Since *f*_2_(*t*)*e*^-λ_3_*t*^ is integrable, the validity of the convergence and expectation follow directly from Theorem 4.7. Lemma 3.5 gives the Laplace transform of *V*_3_.

#### Lemma A.1.

*For i* ≤ 3, 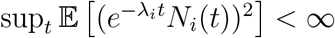.

*Proof.* By Lemma 5 in [10], we know that inductively, if 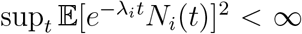 and λ_*i*_ < λ_*i*+1_ holds, then 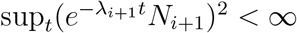. In this case, since 0 = λ_0_ = λ_1_ < λ_2_ < λ_3_, we only need to show that

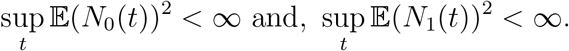

Note that by our transition scheme, one can have

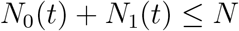

Therefore

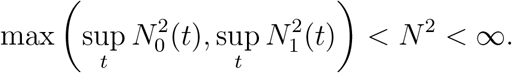

#### Lemma A.2.

*For i* ∈ {2, 3}, 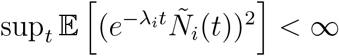.

*Proof.* For *i* = 2, recall that 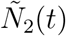 is the second type in a two-type branching process where the first type is 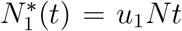. And 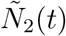 is produced at rate 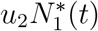. By manipulating the master equation of this two-type process, we obtain the following differential equation of 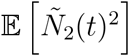,

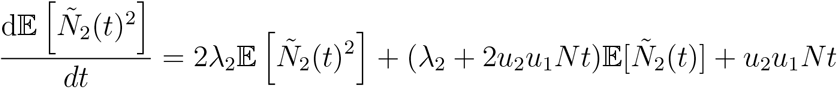

subject to 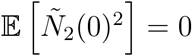. The solution is

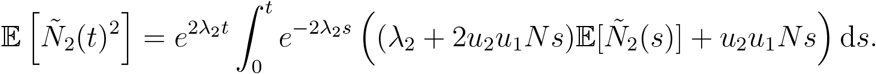

Note that

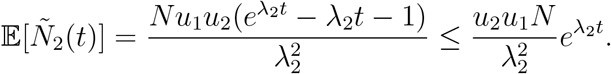

Thus,

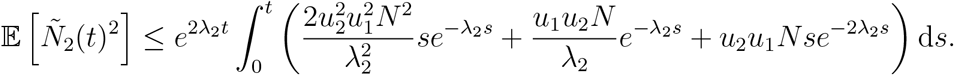

Using the fact that for 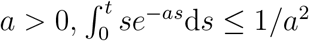 and 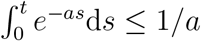, we get 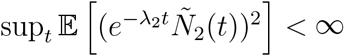.

For 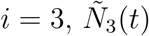 is produced at rate 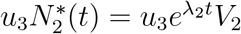. By the master equation we get the following differential equation:

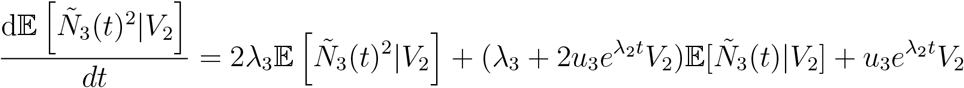

subject to 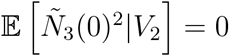. The solution is

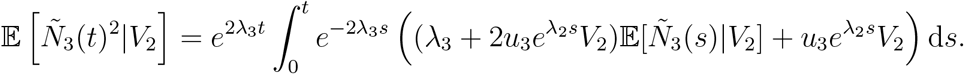

Note that

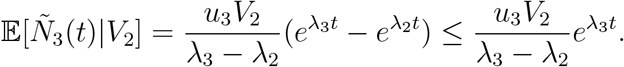

Thus we have

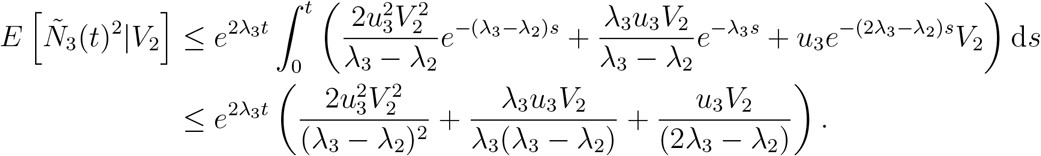

By Lemma 3.5, one can compute the moments of *V*_2_ by it’s Laplace transform. Then we get

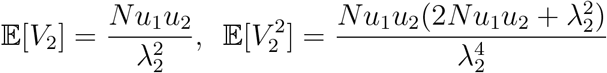

Hence we conclude that 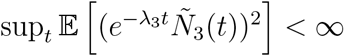.

## B. Closed from formulas of analytic distributions

In the main text, we have omitted a few cumbersome formulas to increase readability. Here we present their closed form expressions. We begin with the analytic probability distribution functions of *τ*_4_.

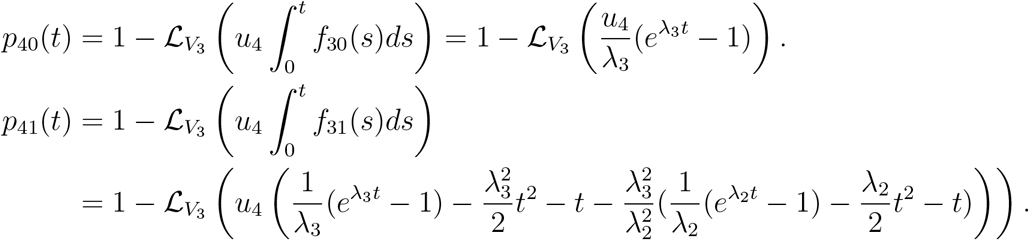

where

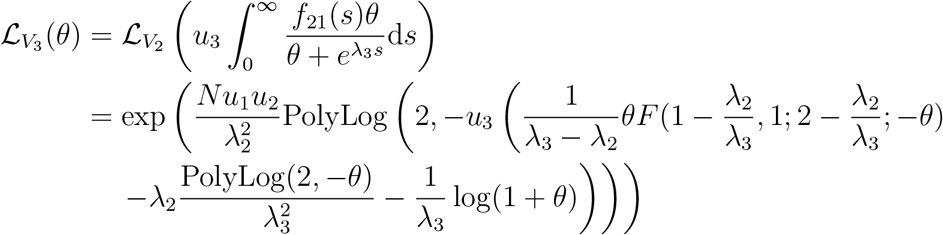

In the equation above, *F*(*a, b; c; z*) is the hypergeometric function.

The two results of skipping type-4 are

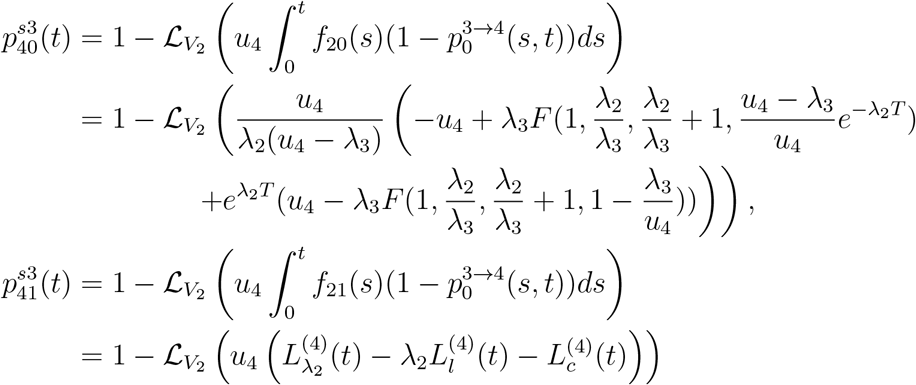

where

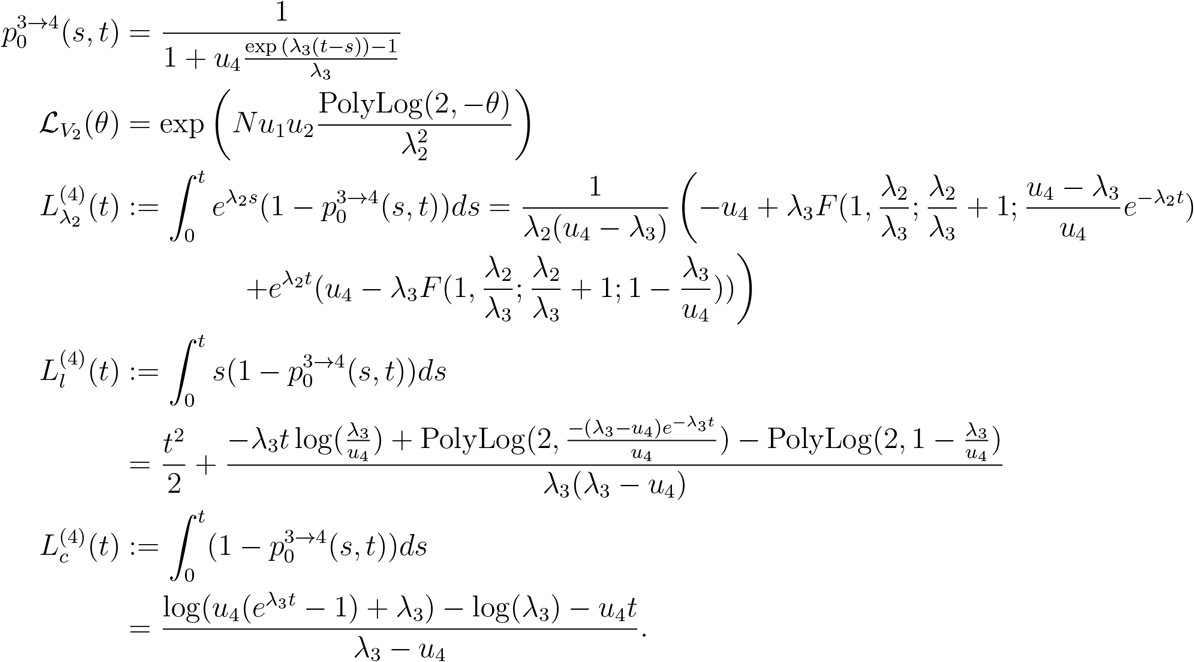

For waiting time distributions of the first type-5 crypt, our estimations are 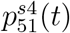 and 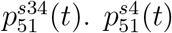 can be expressed explicitly.

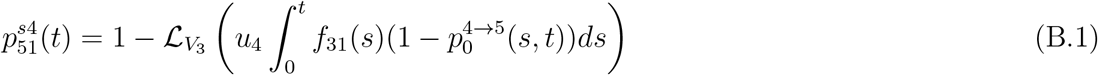

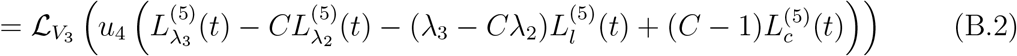

where

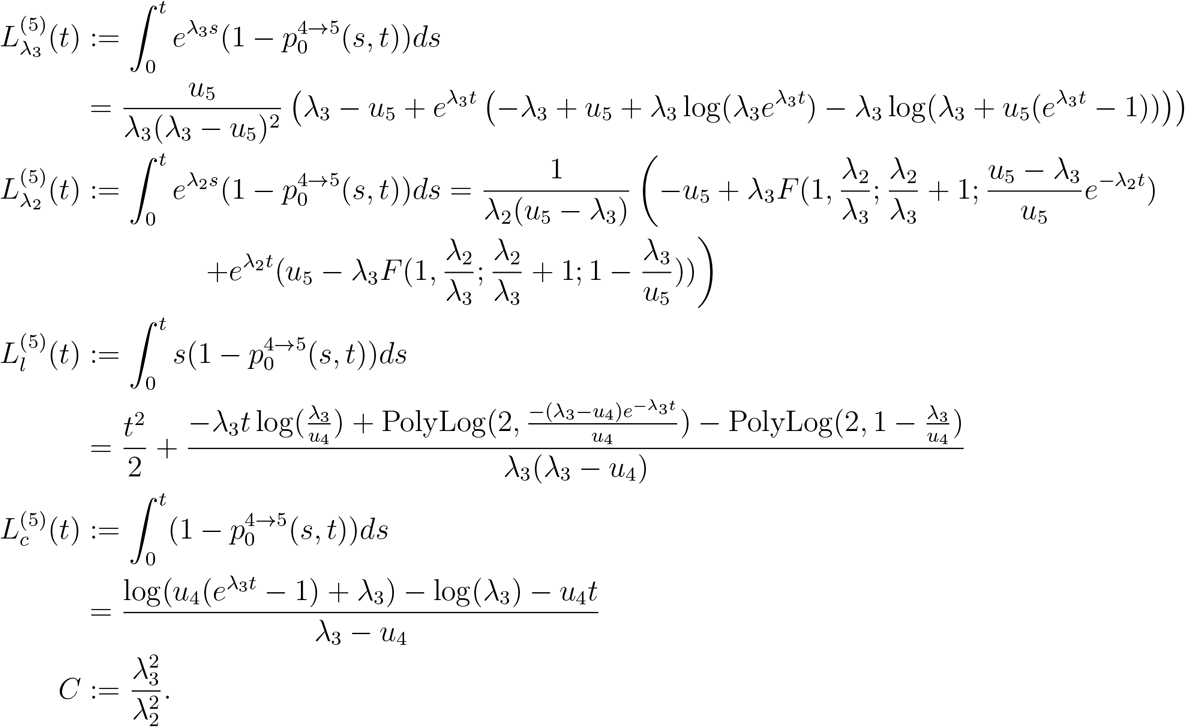

Define *I*(*t*) =

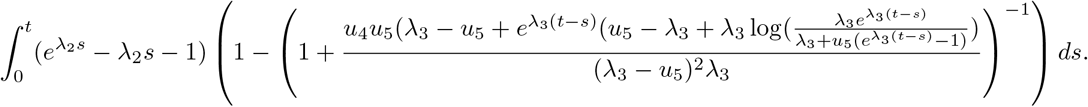

Then we can have

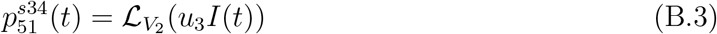

Unfortunately, we didn’t find a way to solve the integral *I*(*t*) explicitly. Nevertheless, we can compute this value numerically.

